# Convergence of glandular trichome morphology and chemistry in two montane monkeyflower species

**DOI:** 10.1101/827220

**Authors:** Sofía Bustamante Eguiguren, Ha An Nguyen, Alexis Caldwell, Kristine A. Nolin, Carrie A. Wu

**Affiliations:** Department of Biology, University of Richmond, 138 UR Drive, Richmond VA, 23173, USA; Department of Chemistry, University of Richmond, 138 UR Drive, Richmond VA, 23173, USA

**Keywords:** Glandular trichome, Secretions, Morphology, Histochemistry, TLC stains, *Mimulus*

## Abstract

Two distantly-related North American montane monkeyflower species, *Mimulus lewisii* and *Mimulus tilingii,* possess glandular trichomes. In this study, we characterized the morphological and histochemical features of these glandular trichomes. For each species, we used traditional light microscopy and scanning electron microscopy (SEM) to examine morphological characteristics, and determined the main components of the secretory products using histochemical and thin layer chromatography (TLC) staining techniques. We identified type VI glandular trichomes on leaf surfaces in both species of monkeyflowers. These trichomes exhibited stalk-cell lengths and head-cell counts that varied across adaxial and abaxial leaf surfaces, stems, and sepals. Both species contained secretory products within the cuticle of the trichome head, which releases the subcuticular metabolites when ruptured. Histochemical tests in both *M. lewisii* and *M. tilingii* confirmed that secretions contained lipids and polysaccharides. TLC plate staining indicated the presence of UV active compounds with polyalcohols, lipids, and amines. The common morphology and chemistry of the glandular trichomes suggests an analogous response to similar environmental conditions in these evolutionary distant montane monkeyflowers.

## 1. Introduction

Trichomes are small hairlike structures that protrude from the epidermis of above-ground vegetative and reproductive tissue (Theobald et al., 1979; Holeski et al., 2013). Leaf trichome morphology, function, and density varies considerably across individuals, populations, and species (Kärkkäinen and Ågren, 2002; Dalin et al., 2008). Nearly one third of all vascular plants contain glandular trichomes, which often cooccur with non-glandular trichomes on the same plant (Holeski et al. 2010; Huchelmann et al. 2017). Glandular trichomes are capable of excreting secondary metabolites that may serve a variety of defensive and physiological roles that contribute to plant fitness (Payne, 1973; Ehleringer, 1984; Wagner, 1991; Agren and Schemske, 1993; Kärkkäinen and Ågren, 2002; Wagner et al., 2004; Schilmiller et al., 2008; Holeski et al., 2010, 2013; Meira et al., 2014; Huchelmann et al., 2017; Tissier et al., 2017; Liu et al., 2019). These secretory structures display high morphological and chemical diversity across the plant kingdom, including variation in the length of the trichome stalk cell relative to the size of the glandular head (Theobald et al., 1979). For example, capitate trichomes have a stalk cell twice as long as their multicellular head, and can serve as physical barriers to increase the leaf boundary layer, thereby regulating leaf temperature and water loss (Theobald et al., 1979; Ehleringer, 1984; Körner, 2003; Glas et al., 2012; Liu et al., 2019). Trichomes may protect tissues from extreme temperatures by reducing heat damage, controlling transpiration, increasing freeze-tolerance, and protecting against damage by UV light (Ehleringer, J., 1984; Gravano et al., 1998; Werker, 2000; Larcher, 2001; Kärkkäinen and Ågren, 2002; Körner, 2003; Wagner et al., 2004; Combrinck et al., 2007; Huttunen et al., 2010; Mershon et al., 2015). Further, glandular trichome secretions can have important roles in pollination, seed dispersion, and inter-plant signaling (Levin, 1973; Wagner, 1991; Holeski et al., 2010; Kärkkäinen and Ågren, 2002; Schilmiller et al., 2008; Holeski et al., 2013; Meira et al., 2014; Tissier et al., 2017; Liu et al., 2019). Some leaf trichomes protect plants from herbivory by secreting fluids that interfere with insect activity (Levin, 1973; Wagner, 1991; Agren and Schemske, 1993; Elle and Hare, 2000; Malakar and Tingey, 2000; Handley et al., 2005; Holeski et al., 2010; Kärkkäinen and Ågren, 2002; Schilmiller et al., 2008; Holeski et al., 2013; Meira et al., 2014; Tissier et al., 2017; Liu et al., 2019). Trichome secretions that possess antifungal, antibiotic, and antithrombotic properties can also defend the plant form pathogens (Dos Santos Tozin and Rodrigues, 2017; Haratym and Weryszko-Chmielewska, 2017; Tissier et al., 2017; Liu et al., 2019). Indeed, secretions from several plant species have been harvested for pharmacological studies as possible alternatives to conventional synthetic antibiotics (Liu et al., 2019). The molecular characteristics of the trichome secretion compounds, such as terpenes and acyl sugars, dictate their function (Gershenzon and Dudareva, 2007; Schilmiller et al., 2008; Huchelmann et al., 2017; Liu et al., 2019).

The genus *Mimulus* (Phrymaceae, updated to *Erythranthe* by Barker et al., 2012; but see Lowry et al. in press) contains 160-200 species that exhibit tremendous phenotypic variation, and has served as a model system for ecological adaptation, speciation, and chromosomal evolution (Wu et al., 2008; Yuan, 2019). In this study, we compare trichome morphology and function of two distantly related species of monkeyflowers, *Mimulus lewisii* (section Erythranthe) and *M. tilingii* (section Simiolus; Beardsley et al., 2004). These two species exhibit substantial range overlap that encompasses montane environments in western North America, likely experience similar environmental conditions, and are characterized by glandular trichomes (Greene, 1895; Schnepf and Busch, 1976; Abrams, 1984; Bohm, 1992; Baldwin et al., 2012.)

Here we performed a comparative analysis of the vegetative glandular trichome morphology and secretion histochemistry of *Mimulus lewisii* and *M. tilingii*. We examined morphology using both light microscopy and scanning electron microscopy (SEM; Ascensão and Pais, 1998; Haratym and Weryszko-Chmielewska, 2017; Rodriguez et al., 2018). We used histochemical tests to elucidate categories of compounds found in the glandular secretions from each species and used TLC to identify the functional groups of the compounds in the secretions. We ask whether the ecological overlap in the range of these two species is reflected in similar trichome structure and function, in spite of their evolutionary distance (Beardsley et al., 2004; Nie et al., 2006).

## 2. Material and Methods

### 2.1. Plant materials

Plants used in this study originated from field-collected lineages centrally-located within the geographic range of each species. Seeds from a *Mimulus lewisii* population in the south-central Oregon, USA, portion of the Modoc Plateau were provided by Paul Beardsley, and belong to the northern race of *M. lewisii* (Heisey et al., 1971; Beardsley et al., 2004; Baldwin et al., 2012). *Mimulus tilingii* seeds were collected from a population in the White Mountains in Inyo County, California, USA (N 37°12.720’ and W 118°36.627’). For both species, we propagated the lineages through self-fertilization for three generations in the greenhouse to reduce maternal effects. Seeds were planted in Fafard 4P potting soil, stratified at 4°C for 7 days, then germinated and maintained in a walk-in custom-built growth chamber (Environmental Growth Chambers, Chagrin Falls, Ohio, USA) under long-day photoperiod conditions (16h light at 22°C/8 h dark at 18°C) and 50% relative humidity. Plants were watered daily to soil saturation, with Peters Professional 20-15-20 fertilizer at 300 ppm added weekly. We sampled vegetative material from leaves, stems, and sepals plants that were in full bloom.

### 2.2. Morphology and distribution of glandular trichomes

To study the morphological features of trichomes in these species, we used multiple complementary microscopy techniques to obtain a thorough characterization of their structure.

#### 2.2.1. Light microscopy

We imaged the glandular trichomes using temporary wet mounts. Fresh leaf tissue was collected from the edge of mature leaves using a razor blade, cut into ca. 5 mm^2^ pieces, and placed on a slide with a droplet of distilled water. Photomicrographs of wet mount leaves were made with a Nikon CoolPix 990 digital camera fitted with a Martin Microscope S/N 0120 adapter on a Nikon Optiphot compound light microscope and Nikon SMZ10 dissecting microscope.

#### 2.2.2. Trichome density

We calculated trichome density on both abaxial and adaxial leaf surfaces for both species from the light micrographs. Trichomes were counted at 4.9X magnification under a dissecting microscope, and trichome density was calculated using FIJI software (Schindelin et al., 2012) on the adaxial (*M. lewisii* n=7; *M. tilingii* n=6) and the abaxial (*M. lewisii* n=5; *M. tilingii* n=6) surfaces. The densities between species and between the leaf surfaces were compared with two-sample t-tests.

#### 2.2.3. Scanning Electron Microscopy (SEM)

To examine the three dimensional structure of trichomes, we captured images using SEM. Following the protocol of Talbot and White (2013), 17 subsections of *Mimulus lewisii* and 11 subsections of *M. tilingii* leaves (5 mm x 5 mm each) were fixed in 100% methanol for 25 min at room temperature. Subsequently, the plant material was dehydrated twice in 100% ethanol for 30 min. Once dehydrated, the samples were critical-point dried in an EMS 850 Critical Point Dryer, and sputter coated with gold-palladium following manufacturer’s protocols for the Denton Desk IV sputter coater. The adaxial and abaxial leaf surfaces were imaged under a JOEL JSM-6360LV scanning electron microscope at 10kV accelerating voltage.

### 2.3. Secretion Chemistry

To characterize chemical characteristics of the trichome glandular secretions, we used histochemical techniques and other staining processes following thin layer chromatography (TLC). The glandular secretions for *Mimulus lewisii* and *M. tilingii* were analyzed with staining techniques both *in vivo* on the trichomes and *in vitro* after isolating the secretions from the leaves.

#### 2.3.1. Histochemistry

The main classes of metabolites present in *Mimulus lewisii* and *M. tilingii* leaf glandular secretions were determined using protocols from Haratym and Weryszko-Chmielewska (2017) for the following histochemical tests: potassium dichromate for tannins (Gabe, 1968), Toluidine Blue O for polysaccharides (Serrato-Valenti et al., 1997), Ruthenium Red for polysaccharides that are not cellulose (Johansen, 1940), Nile Blue for acidic and neutral lipids (Jensen, 1962), Sudan Black B and Sudan III for lipids (Johansen, 1940; Lison, 1960), and Neutral Red for essential oils and lipids (Clark, 1981). All stains were matched with an unstained control. We imaged freshly stained tissue following the light microscopy techniques described previously (Section 2.2.1) on a Nikon Eclipse E600 microscope.

#### 2.3.2. Extraction and isolation of secretions

*Mimulus lewisii* and *M. tilingii* trichome secretions were isolated from plant tissue using 95% ethanol leaf washes. Leaf washes allow isolation of the trichome secretions without disrupting the leaf integrity (Asai et al., 2012). After optimizing the method from Asai et al. (2012), each freshly-collected leaf was rinsed for 15 seconds. To obtain sufficient material for subsequent analyses, 699 individual leaf washes of *Mimulus lewisii* were pooled together, as well as 500 leaf washes for *M. tilingii*. After removing the solvent with a Buchi Rotavapor R-210, secretions were resuspended in ethyl acetate. To screen for functional groups, we spotted the secretions on thin layer chromatography (TLC) silica plates prior to exposure to the staining solutions. Plates were visualized following standard procedures from Tuchstone (1992), including: UV light only for conjugated pi-systems; vanillin-sulfuric acid for alcohols, ketones, bile acids, and steroids; phosphomolybdic acid for steroids and lipids; cerium ammonium molybdate for polyalcohols; potassium permanganate for alkenes, alkynes, alcohols, and amines; p-anisaldehyde for polysaccharides; ninhydrin for amines; diphenylamine in EtOH for nitrate esters; copper(II) sulfate for sulfur containing glycosides; and 2, 4-dinitrophenylhydrazine for aldehydes and ketones.

## 3. Results

### 3.1. Morphology and distribution of glandular trichomes

Both *Mimulus lewisii* and *M. tilingii* leaves possess predominantly capitate glandular trichomes on the adaxial and abaxial surfaces (Figs. 1 and 2).

**Fig. 1.**
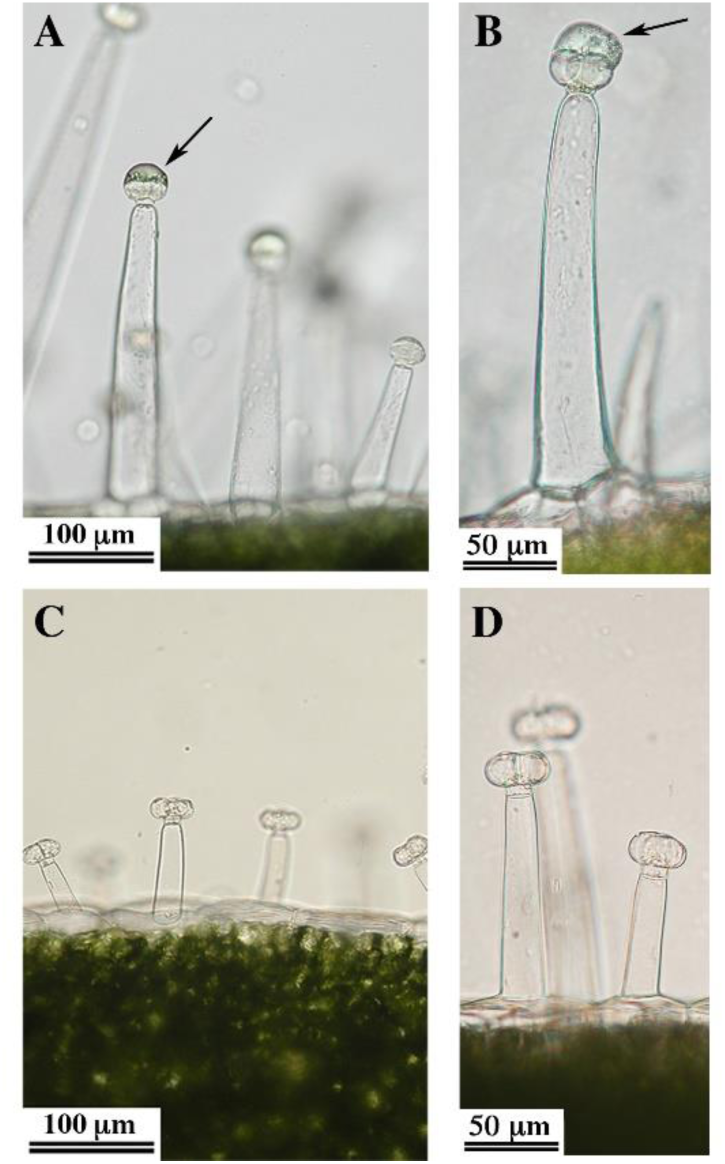
Long-stalk-multicelled glandular trichomes found on leaf surfaces of both *Mimulus lewisii* and *M. tilingii* visualized by light microscopy. The stalk cell length varies in *M. lewisii* (**A, B**) trichomes, but remains relatively constant in *M. tilingii* (**C, D**) trichomes. Trichomes on both species have a single-celled stalk, a neck cell, and a multicellular head that is surrounded by the secretions, corresponding to type VI trichome. The glandular secretions (**A,B**) the head of *M. lewisii* trichomes are indicated by arrows. Note the scalebars.

**Fig. 2.**
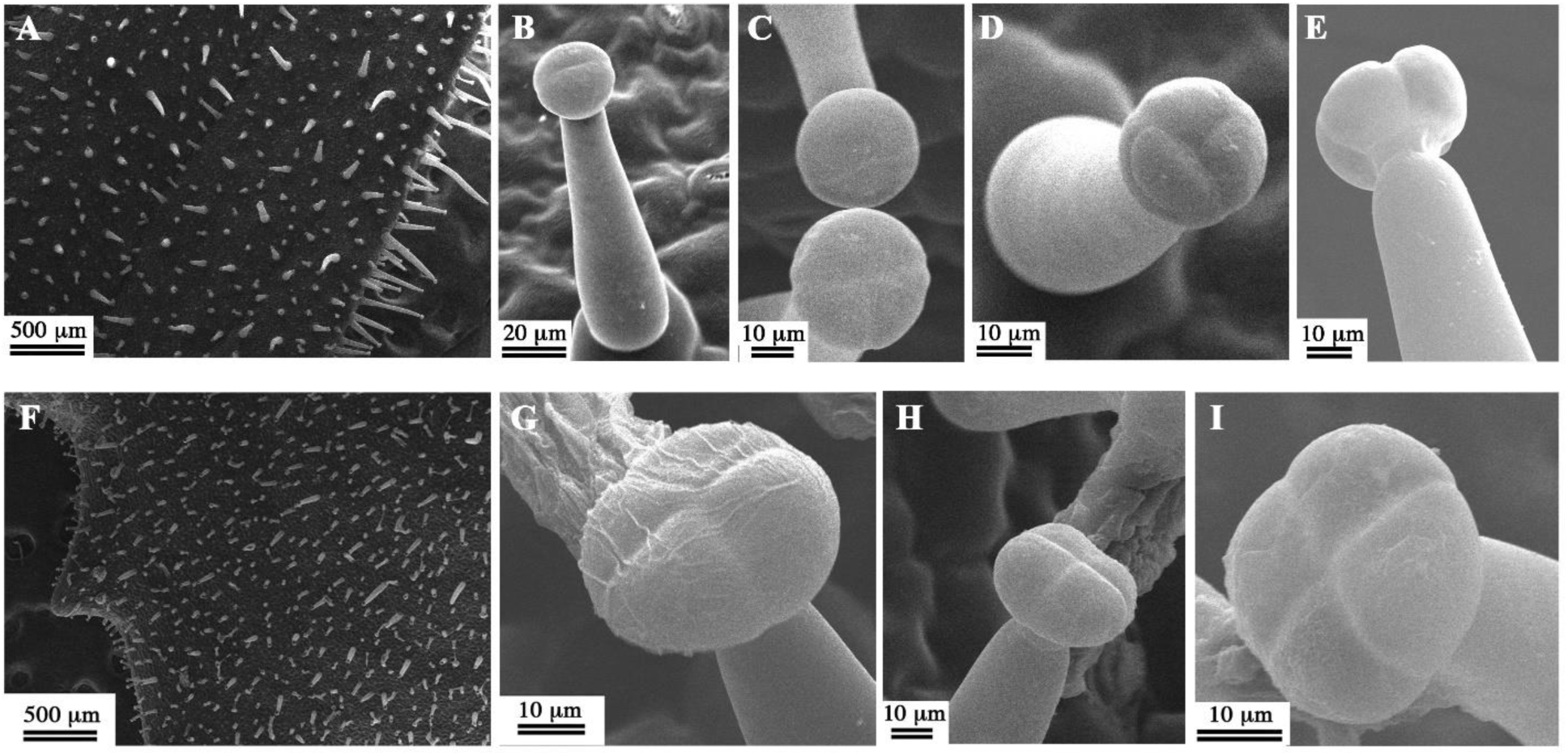
Morphology of *Mimulus lewisii* (**A-E**) and *M. tilingii* (**F-I**) glandular trichomes visualized by SEM consistent with type VI capitate glandular trichomes that have a single-cell stalk, a neck cell, and multicellular heads. Both, *M. lewisii* (**A**) and *M. tilingii* (**F**), have evenly distributed trichomes in adaxial (shown) and abaxial (not shown) leaf surfaces. The stalk cells are the main source of height variability among all the trichomes in *Mimulus*. Although bicellular trichome heads (**B-C**) are the most common type VI trichome structure present in *M. lewisii*, tricellular (**C-D**) and tetracellular glandular heads (**E**) were also observed in a single plane. The glandular trichomes of *M. tilingii* had only tetracellular glandular heads (**G-I**) in a similar planar structure. The secretions of *M. tilingii* trichomes are visible after the cuticle has been removed (**G**). Note scalebar differences among panels.

#### 3.1.1. Trichome density and relative size

In both species, trichomes were distributed regularly across the leaf surface (Figs. 2A and 3A). However, trichome density was nearly twice as high in *Mimulus tilingii* (mean ± SE 35.5 ± 2.30 trichomes per mm^2^) than in *M. lewisii* (17.4 ± 1.17 trichomes per mm^2^; two-sample t-test, n = 24, df = 22, t=-6.99, p < 0.001), with no significant difference between abaxial and adaxial leaf surfaces within a given species (Table 1; two-sample t-test, each n=12, df=10, *M. lewisii* contrast t= 0.81and *M. tilingii* contrast t= −0.37; both p > 0.05). Qualitatively, the trichomes appeared overall longer, though more variable in length, in *M. lewisii* than in *M. tilingii* (Fig. 1).

**Table 1.** Average trichome densities (mean ± SEM trichome/mm^2^, n=12 leaves per species) of mature *Mimulus* leaf surfaces.

| Leaf surface | *M. lewisii* | *M. tilingii* |
| --- | --- | --- |
| Adaxial | 17.1 ± 1.87 | 36.34 ± 2.60 |
| Abaxial | 17.7 ± 1.30 | 34.6 ± 4.04 |
| Average across surfaces | 17.4 ± 1.17 | 35.5 ± 2.30 |

#### 3.1.2. Trichome types

The glandular trichomes on *Mimulus lewisii* and *M. tilingii* show extensive morphological similarity (Fig. 1). Leaves from both species contain capitate trichomes with a single cell stalk, a unicellular neck, and a multicellular head that is surrounded by a cuticle containing the secretory compounds (Figs. 1 and 2). Trichomes with these characteristics are generally classified at type VI (Luckwill, 1943; Channarayappa et al., 1992; Glas et al., 2012).

*Mimulus lewisii* glandular trichomes are long projecting structures from the epidermal surfaces, with consistent density and morphology across all the tissue examined. In SEM micrographs of *M. lewisii*, the whole trichome surfaces have smooth surfaces (Fig. 2B-E). Most trichomes had a single basal cell, although occasionally larger trichomes appeared to have more than one basal cell (Fig. 1B). In all samples analyzed, the stalks and the necks were single cells (Figs. 1B, C and 2B, D, E). Within each species, the length of the stalk cell was the most variable component of the trichomes (Figs. 1B, C and 2A, F). The heads contained 2-4 cells arranged in a single plane, which could be observed through SEM only after the cuticle was removed in the fixation process (Fig. 2B-E). We could also observe the multicellular heads with the compound light microscope, which revealed the cuticle with the secretions above or surrounding the cells (Fig. 1B,C).

*Mimulus tilingii* glandular trichomes also have consistent density and morphology across the leaf surface, although they appeared generally shorter than *M. lewisii* trichomes. SEM micrographs showed trichomes that were minutely verrucose and, like *M. lewisii*, had a single basal cell (Fig. 2G-I). As in *M. lewisii*, the *M. tilingii* trichomes varied in height primarily due to varied lengths of the stalk cells (Figs. 1C and 2F). *Mimulus tilingii* trichomes also had a single neck cell that was thinner than the head and the stalk (Figs. 1C-D and 2H). We consistently observed tetracellular heads in the glandular trichomes of *M. tilingii* (Fig. 2G-I). In some of the SEM images, the secretions were still visible on the trichome heads despite the harsh dehydration process (Fig. 2G-I). Some light microscopy images showed the cuticle with the subcuticular secretions, consistent with secretions being released following cuticle rupture (Fig. 3P).

**Fig. 3.**
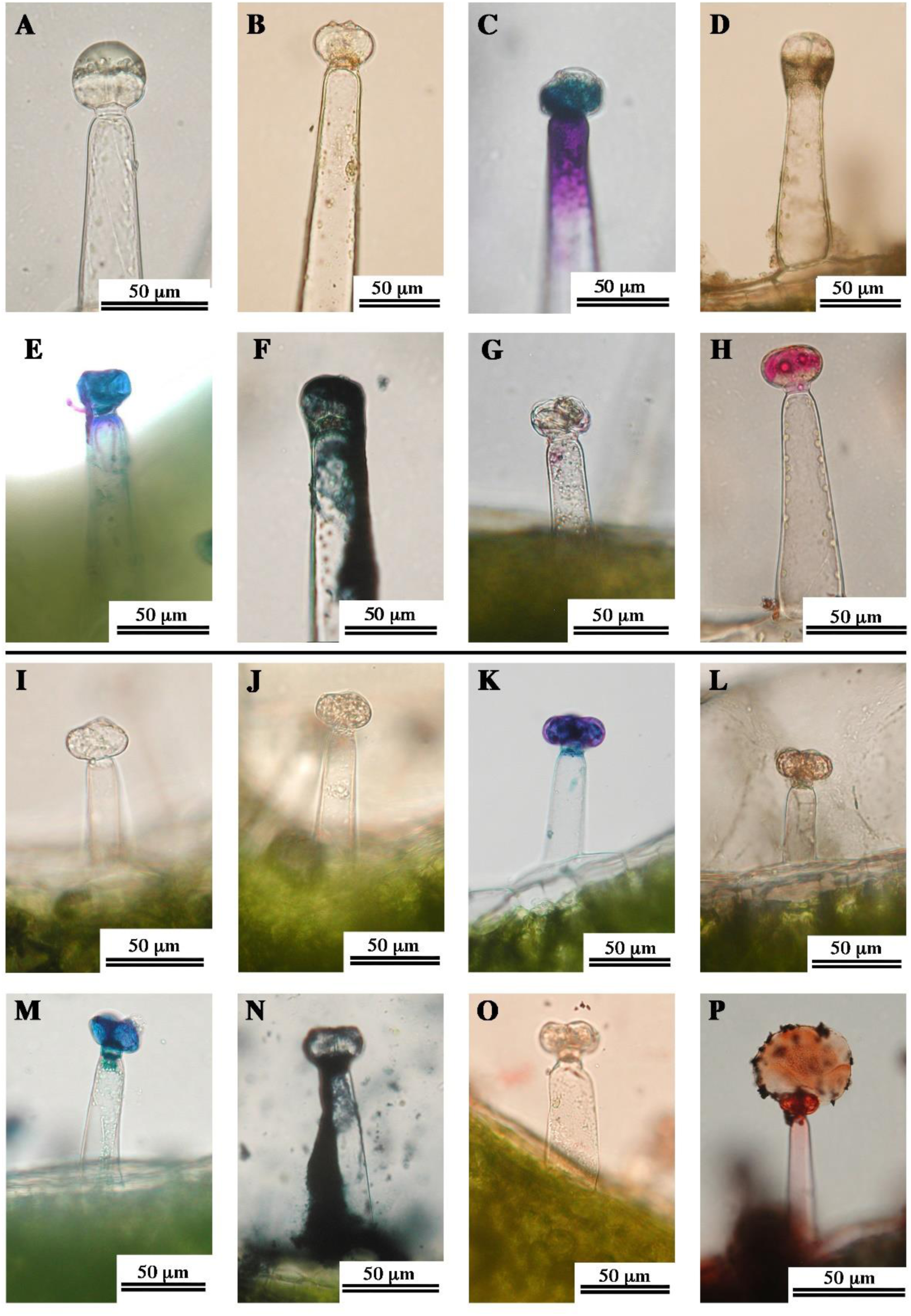
Histochemical characterization of secretions of *Mimulus lewisii* (**A-H**) and *M. tilingii* (**I-P**) visualized by light microscopy. The main classes of metabolites in the resins of both species were investigated using fresh leaf sections with seven histochemical tests, using protocols modified from Haratym and Weryszko-Chmielewska (2017). Unstained *M. lewisii* (**A**) and *M. tilingii* (**I**) capitate trichomes, with visible secretory glandular bicellular head, neck cell, and stalk cell. Staining with potassium chromate indicate a lack of tannins in trichomes of *M. lewisii* (**B**) and *M. tilingii* (**J**). Polysaccharides stained with Toluidine blue in the stalk cell and head cell of *M. lewisii* (**C**) and only the head cells of *M. tilingii* (**K**). Low relative abundance of polysaccharides stained with Ruthenium red was seen in *M. lewisii* (**D**) and *M. tilingii* (**L**) trichomes. Neutral lipids stained with Nile blue in the glandular head of *M. lewisii* (**E**) and *M. tilingii* (**M**). There is an abundant lipidic resin stained with Sudan Black B on the secretions of*M. lewisii* (**F**) and *M. tilingii* (**N**). Lower concentration of lipids in the resin stained with Sudan III in *M. lewisii* (**G**) and *M. tilingii* (**O**). Lipids stained with Neutral red in the glandular head of *M. lewisii* (**H**) and the subcuticular area and head cells of *M. tilingii* (**P**).

### 3.2. Histochemical analysis

#### 3.2.1. *In vivo* analysis

Histochemical staining revealed numerous substances in trichome secretions of both *Mimulus lewisii* and *M. tilingii* (Table 2). Fresh unstained sections appeared transparent both species (Fig. 3A, I). Trichome secretions stained positively for polysaccharides and lipids, but did not stain for tannins (Fig. 3). No substantial staining of tannins was observed in the secretions with potassium dichromate treatment, but the cell walls appear darker (Fig. 3B, J). A high polysaccharide concentration was visible in the head cells of both with Toluidine Blue O (Fig. 3C, K), and was also observed in the stalk cell of *M. lewisii* (Fig. 3C). Ruthenium Red treatment also indicated polysaccharides in the secretions of the two species (Fig. 3D, L). Both species were positive for acid lipids based on staining with Nile Blue (Fig. 3E, M), and positive for lipids when stained with Sudan Black B, Sudan III, and Neutral Red (Fig. 3F-H, N-P). When treated with Sudan III, only small lipidic vesicles were visible in both species (Fig. 3G, O). While Neutral Red stains the head cells of both species, *M. tilingii* shows higher lipidic concentrations in the surrounding secretions based on stain intensity (Fig. 3H, P).

**Table 2.** Histochemical identification of compounds in the capitate type VI trichomes of *Mimulus lewisii* and *M. tilingii*. Staining responses summarized from Haratym and Weryszko-Chmielewska (2017). Representative images of staining results are shown in Figure 1.

| Stain | Compound | Color observed | Target response |  |
| --- | --- | --- | --- | --- |
|  |  |  | *M. lewisii* | *M. tilingii* |
| Potassium dichromate | Tannins | Brown | - | - |
| Toluidine Blue O | Polysaccharides | Blue/purple | + | + |
| Ruthenium Red | Polysaccharides | Crimson | (+) | (+) |
| Nile Blue | Acid lipids | Blue | + | + |
| Sudan Black B | Lipids | Dark blue | + | + |
| Sudan III | Lipids | Orange | - | - |
| Neutral Red | Lipids | Red | + | + |

'-' negative response, '(+)' subtle response, '+' positive response

#### 3.2.2. Thin layer chromatography

*Mimulus lewisii* and *M. tilingii* leaf glandular secretions responded positively to all tests for functional groups within the compounds we examined (Table 3). When *M. lewisii* and *M. tilingii* glandular secretions were spotted on TLC plates and exposed to short-wave UV light, the spots from both species fluoresced, indicating the presence of conjugated pi systems in the compounds. Vanillin-sulfuric acid tested positive for alcohols, ketones, bile acids or steroids for both species, though coloration was darker in *M. lewisii*. Steroids, lipids and antioxidants were also present in the secretory compounds, as evidenced by spots in phosphomolybdic acid. Polyalcohols were positively characterized with cerium ammonium molybdate. Potassium permanganate stained the glandular secretions from both species for alkenes, alkynes, alcohols, and amines, suggesting the presence of double bonded components. The stain for sugars, p-anisaldehyde, were darkly spotted. Staining with ninhydrin indicated low amounts of amines in the secretions for both *M. lewisii* and *M. tilingii*. Nitrate esters were also weakly detected as indicated by yellow spots on diphenylamine in EtOH. Copper (II) sulfate lightly stained for sulfur containing glycosides. Finally, the orange spots on the 2, 4-dinitrophenylhydrazine stained plate confirmed the presence of aldehydes and ketones. Overall, the compounds of both species appear to contain alcohols, lipids, alkynes, sugars, amines, nitrate esters, and some sulfur-containing glycosides.

**Table 3.**
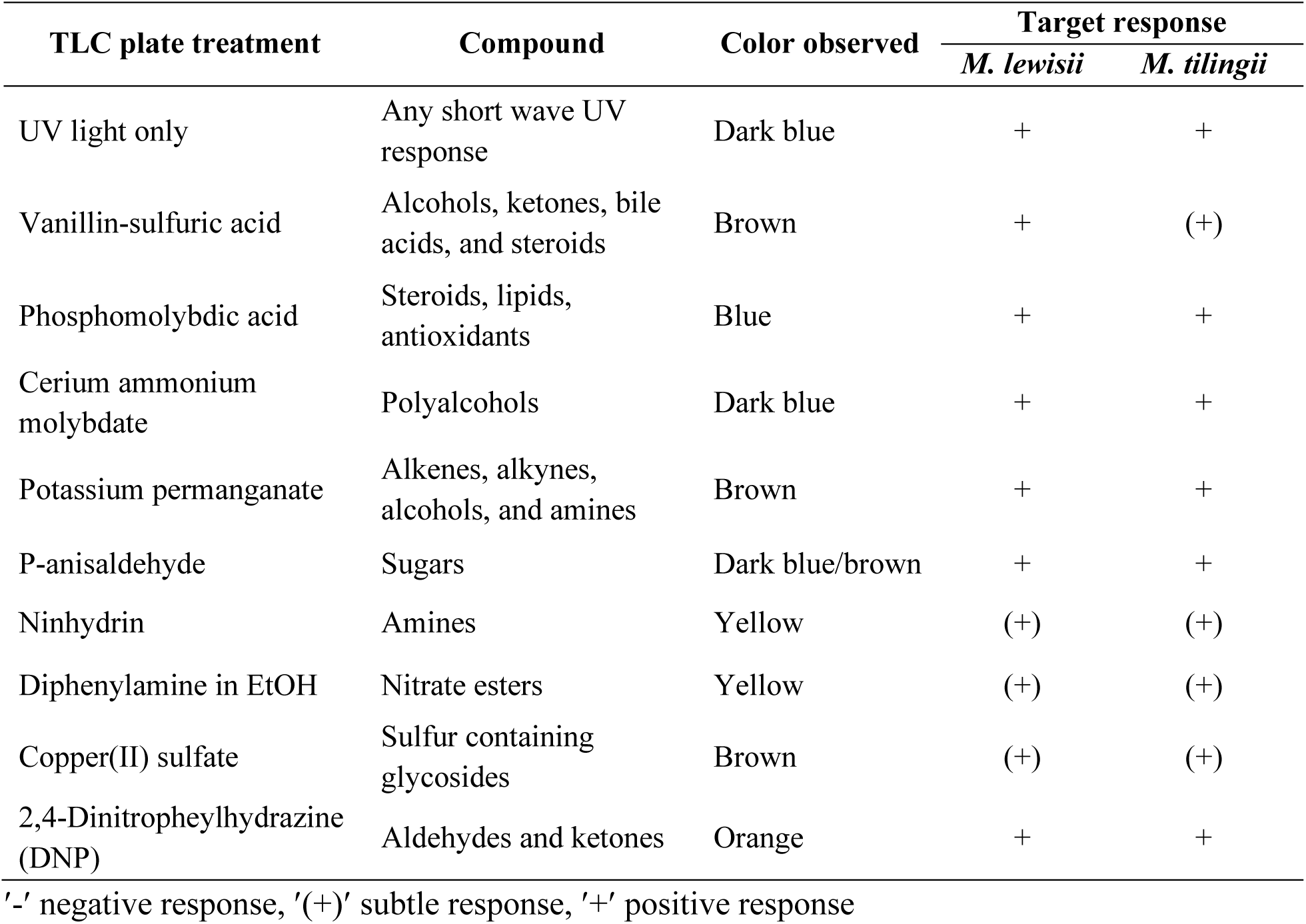
Functional group identification in compounds from the extracted secretions of *Mimulus lewisii* and *M. tilingii* leaves using TLC plate staining, following methods from Tuchstone (1992).

## 4. Discussion

We determined that two distantly-related montane monkeyflower species, *Mimulus lewisii* and *M. tilingii,* both contain type VI glandular trichomes on vegetative tissues that are characterized by a stalk cell, a neck cell and a multicellular head that produce lipids and polysaccharides. This structure is consistent with the only prior morphological characterization in *M. tilingii* of which we are aware (Schnepf and Busch, 1976) that identified a tetracellular trichome head using light microscopy, and is similar to the straight, unicellular trichomes described in the closely related species *M. guttatus* (Holeski 2007; Holeski et al. 2010).

Capitate trichome morphologies have been classified into eight categories, largely based on the number of cells, length, and shape (Luckwill, 1943; Channarayappa et al., 1992; Glas et al., 2012). The trichome morphology of both *Mimulus* species studied here are consistent with type VI, which have a single stalk cell, a neck cell, and a multicellular head within the same plane (Luckwill, 1943; Channarayappa et al., 1992; Glas et al., 2012). Most trichomes observed in *M. lewisii* are bicellular or tricellular in the glandular head, while *M. tilingii* have primarily tetracellular heads, consistent with descriptions by Schepf and Busch (1976). Other montane species, *Solanum lycoperiscum* and *S. tuberosum,* also have type VI glandular trichomes with tetracellular heads (Kang et al., 2010; Bergau et al., 2015; Cho et al., 2017). The presence of a cuticle protecting the secretions in the subcuticular space of the trichomes that we observed in *M. lewisii* and *M. tilingii* was also identified in *S. lycoperiscum* and *S. tuberosum*, suggesting this structure could function to sequester and store the secretions until a physical disturbance, such as water droplets or insect visitation, causes the cuticle to rupture (Tissier et al., 2017). This extracellular structure thus may prevent intracellular accumulation and self-toxicity (Tissier et al., 2017).

Trichome length, size, and density have been found to vary in response to the environment factors in dicotyledonous angiosperms (Theobald et al., 1979; Malakar and Tingey, 2003; Holeski, 2007; Holeski et al. 2010). In species such as *Potentilla glandulosa* growing in the Sierra Nevada mountains, trichome density decreases with altitude, which was attributed to responses to herbivory and oviposition, rather than elevational factors (Levin, 1973). In other systems, however, decreased trichome density with altitude has been proposed as a response to a reduced number of herbivores present in the higher ranges, such that the resources can be allocated elsewhere (Wilkens et al., 1996; Kofidis and Bosabaldis, 2008; Horgan et al., 2009). In contrast, trichome density has been reported to increase with altitude in potatoes (Horgan et al., 2009), tomatoes (Wilkens et al., 1996), and salva-de-marajó (Tozin et al., 2015). Among perennial coastal *M. guttatus* populations, average trichome density also varies with elevation. Holeski (2007) suggested density changes inversely with elevation as a response to herbivory, and also shows phenotypic plasticity. It is possible that, like *Mimulus guttatus, M. lewisii* and *M. tilingii* may exhibit plasticity in trichome density, largely in response to herbivory (Holeski, 2007; Holeski et al., 2010), although future studies are needed. Additionally, although our study did not include density measurements between populations, we suspect that the trichome density in *M. lewisii* and *M. tilingii* could have other functionalities beyond herbivore defense.

Trichomes tend to be precocious structures that develop before the leaf fully matures (Rodriguez et al., 2018). As leaf maturation proceeds, intercalary growth increases the average distance between trichomes, such that greater intercalary growth produces a larger leaf with reduced trichome density (Rodriguez et al., 2018). We hypothesize that mature leaves of *M. lewisii,* which are larger than those of *M. tilingii*, may have lower trichome density in part because of such greater intercalary growth. Interestingly, we found that the trichome density did not differ significantly between the adaxial and abaxial surfaces within each species, yet other studies have reported density variation between the opposing leaf surfaces in other species (Bergau et al., 2015; Rodriguez et al., 2018). For example, in the fern *Notholanea sulphurea*, trichomes in the adaxial surface are only present in younger plants, purportedly for protection during early developmental stages (Ascensão et al., 1995; Werker, 2000; Rodriguez et al., 2018).

Trichome secretions of both monkeyflower species contained polysaccharides and lipids, but no tannins. Polysaccharides are common secretory compounds for external defense in several other species, including *Marriubium vulgare* and *Notholaena sulphurea* (Schmilmiller et al., 2008; Keefover-Ring et al., 2014; Haratym and Weryszko-Chmielewska, 2017; Rodriguez et al., 2018; Liu et al., 2019). Additionally, the presence of lipophilic compounds has been widely described in glandular secretions of numerous species, especially terpenoids and flavonoids (Asai et al., 2012; Liu et al., 2019). For instance, terpenoids are a common lipidic compound category, which are biosynthetically derived from five-carbon rings, such as salvorin A and (-)-menthol (Liu et al., 2019). Additionally, flavonoids are also commonly present in glandular trichomes secretions across numerous species (Wollenweber and Schneider, 2000; Kang et al., 2010; Haratym and Weryszko-Chmielewska, 2017; Rodriguez et al., 2018; Liu et al., 2019). Bohm (1992) characterized the secondary chemistry of the entire leaf in *M. lewisii* and described several flavonoids, but these were not present in our analysis of the trichome secretions of either *Mimulus* species. Because we instead specifically isolated external trichome secretions, this divergence in flavonoid detection between our study and that of Bohn (1992) suggests separate metabolic pathways or metabolic packaging in the production of internal compounds and those that are excreted through the trichomes.

Polysaccharides are present in the secretory products from both *Mimulus* species. Consistent with our positive Toluidine Blue O staining, Schnepf and Busch (1976) hypothesized that the secretions of *M. tilingii* contain carbohydrates based on their observations of Golgi bodies in the trichome head cells. Interestingly, the polysaccharides in the secretions of *M. lewisii* were present in the head cell as well as the stalk cell, which suggests certain compounds are produced in the Golgi of stalk cells, then transported to the multicellular head prior to secretion. It is possible that the polysaccharides stained are precursors of lipidic compounds, which have been suggested to play a role in environmental stress response (Schilmiller et al., 2008). The high concentrations of lipids in the glandular heads of both *M. lewisii* and *M. tilingii* suggests that they are produced in the trichome head primarily for secretion, as compounds produced in the trichomes are generally not transported back to the rest of the plant because they can often be hazardous to its internal metabolism (Schilmiller et al., 2008; Tissier et al., 2017). Furthermore, the TLC plate spotting of the isolated glandular secretions revealed a variety of functional groups present in both species including alcohols, lipids, alkynes, sugars, amines, nitrate esters, and some sulfur-containing glycosides. Many studies have found these functional groups in terpenes, a common natural product biosynthesized from five-carbon compounds (Gershenzon and Dudareva, 2007; Huchelmann et al., 2017; Liu et al., 2019). Secretions from both species have conjugated pi systems which also correspond with terpenoids (Schilmiller et al., 2008). Terpenes are commonly synthesized by similar capitate trichomes across several species such as tomatoes (Schilmiller et al., 2008). Additional characterization of the molecular characteristics of the compounds in the secretions may further our understanding their functional role, such as freeze tolerance or protection against other stresses (Gershenzon and Dudareva, 2007; Schilmiller et al., 2008; Huchelmann et al., 2017; Liu et al., 2019). Other studies have found that trichome secretions can serve as pathogen defenses based on antifungal, antibiotic, and antithrombotic properties (Dos Santos Tozin and Rodrigues, 2017; Haratym and Weryszko-Chmielewska, 2017; Tissier et al., 2017; Liu et al., 2019), or provide protection against UV light and other abiotic stresses (Liu et al., 2019). Additional analyses are needed to determine the specific functional role of these compounds in *M. tilingii* and *M. lewisii*.

The similarities between the species studied here include the trichome structural type, general morphology, and secretion chemistry. *Mimulus lewisii* and *M. tilingii* are exposed to similar environmental stresses found in montane environments, such as below-freezing temperatures and high UV light (Körner, 2003; Wu et al., 2008; Baldwin et al., 2012). Therefore, these trichomes might serve as a physical barrier to prevent intracellular ice formation by creating an air space between the trichome heads and the epidermis, forming an insulation layer to protect the leaves (Azocar et al., 1988; Zhen and Ungerer, 2008; Li et al., 2018). In addition, the lipids produced by these glandular trichomes may serve as a hydrophobic layer that reduces the accumulation of water on the adaxial leaf surface, further limiting freezing within the epidermal tissue. Minimizing the damage from cold temperatures that may occur late in the spring or early in the fall could potentially extend the reproductive period (Körner, 2003). Since *M. lewisii* and *M. tilingii* have similar trichome structures and chemistry, yet belong to different species complexes within the genus (Beardsley et al., 2004), these trichome characteristics could reflect convergent evolution in response to common environmental pressures.

## 5. Conclusions

The results of this work suggest that trichomes of the montane species *Mimulus lewisii* and *M. tilingii* may have converged to a similar form and function in response to shared environmental conditions that characterize their natural range across western North America. Both species have the same type VI glandular trichomes and almost no non-glandular leaf trichomes. The main components of the secretory products from both species were identified as lipids and polysaccharides, with possibly additional terpenoid compounds. A study of the environmental factors these species face will be necessary to determine the functional role of these convergent structures.

## Acknowledgements

This work was supported by funds from the University of Richmond, Richmond, VA. We thank W. John Hayden for helpful comments on an early draft of the manuscript and assistance with microscopy. We also thank Christine Lacy for her assistance with microscopy.

